# Real-time, functional intra-operative localization of rat cavernous nerve network using near-infrared cyanine voltage-sensitive dye imaging

**DOI:** 10.1101/537274

**Authors:** Jeeun Kang, Hanh N. D. Le, Serkan Karakus, Adarsha P. Malla, Maged M. Harraz, Jin U. Kang, Arthur L. Burnett, Emad M. Boctor

## Abstract

Despite current progress achieved in the surgical technique of radical prostatectomy, post-operative complications such as erectile dysfunction and urinary incontinence persist at high incidence rates. In this paper, we present a methodology for functional intra-operative localization of the cavernous nerve (CN) network for nerve-sparing radical prostatectomy using near-infrared cyanine voltage-sensitive dye (VSD) imaging, which visualizes membrane potential variations in the CN and its branches (CNB) in real time. As a proof-of-concept experiment, we demonstrate a functioning complex nerve network in response to electrical stimulation of the CN, which was clearly differentiated from surrounding tissues in an *in vivo* rat prostate model. Stimulation of an erection was confirmed by correlative intracavernosal pressure (ICP) monitoring. Within 10 minutes, we performed trans-fascial staining of the CN by direct VSD administration. Our findings suggest the applicability of VSD imaging for real-time, functional imaging guidance during nerve-sparing radical prostatectomy.

## Introduction

Despite the remarkable evolution in radical prostatectomy (RP) from the radical perineal prostatectomy reported by Young to the nerve-sparing radical retropubic prostatectomy (RRP) described by Walsh^1–3^, post-operative complications such as erectile dysfunction, in particular, persist, partly due to surgical damage to erection producing cavernous nerves (CN) resulting from this prostate cancer treatment. Because of the proximity of the CN to the prostate gland, at an average distance of 2.8 mm, these nerves are at risk for injury during the surgical procedure. Also, it is still unknown to what extent CN branches (CNB), surrounding the prostate with a spray-like distribution^4^, contribute to erectile function. The CNB are contained within the periprostatic and levator fascias, part of which requires dissection to surgically access the prostate. The individual CN are a few hundreds of micrometers in diameter and the CNB are even thinner, making it difficult to predict the exact locations and paths of these nerves, due to inter-patient variability.^5^ Despite the innovation of laparoscopic/robotic RP^6^, post-operative erectile function recovery outcomes have not necessarily improved over time with only approximately 70% resulting in fully restored potency rates at 12 months after surgery.^7^ Irrespective of surgical approach, RP demands an improved understanding of the anatomical features of the target tissue, particularly with respect to the functional anatomy of the CN network for its maximal preservation.^8,9^

Fluorescence (FL) imaging has been advanced as a vital tool in the field of intra-operative imaging. Intriguing commercial solutions have been recently introduced such as the FireFly™ (Intuitive Surgical Inc., United States) and the FLARE™ systems (Curadel ResVet Imaging, LLC., United States)^10,11^, and several exogenous fluorophores targeted to specific tissue types have also been proposed.^12–14^ Enthusiasm for FL imaging in academia and industry originated from its wide field-of-view (FOV) directly superimposable on the surgeon’s visual field, making it amenable to conventional surgeries including RP. Most of the current nerve-specific fluorophores demonstrate nerve networks with high affinity, but this mechanism will only yield stationary locations of nerves, rather than portray their electrophysiological activity based on erection functionality.^13,15^ Furthermore, most of these nerve-specific fluorophores provide a superficial imaging depth because they have peak absorption and emission wavelengths at the visible wavelength range (400-650nm). Although several research presentations have recently suggested the applicability of infrared dyes to intra-operative nerve localization, they are still based on a nerve-specific affinity mechanism, rather than on the functional resolution of a patient’s erectile function.^16,17^ Additionally, nerve labeling with this affinity-based mechanism takes a long time, from a few hours to days. A direct administration method for near-infrared dye was recently proposed to address this problem, but it is still based on the affinity-based mechanism.^14^

Recently, we proposed the novel mechanism of cyanine voltage-sensitive dye (VSD) imaging, which redistributes the concentration of dyes according to cell membrane potential, displaying functional contrast that corresponds to electrophysiological events in biological tissue.^18,19^ In detail, the cyanine dye with positively-charged chemical structure (e.g., IR780 perchlorate) is attracted into the cell membranes when the neuronal cells are in their polarized resting state (i.e., −90mV).^20^ The high VSD concentration inside the cell membrane triggers their aggregation, leading to FL quenching that dissipates the absorbed light energy in the form of thermal energy. Conversely, when the neuronal cells are in their depolarized state, the VSD aggregate is disassembled and dispersed back into the intra-/extra-cellular spaces. This redistribution restores FL emission. This functional contrast change is activated in sequences of neuronal depolarization and repolarization events, and can be quantified by FL imaging at its near-infrared peak absorption and emission wavelengths at 790nm and 820nm, respectively, which is beneficial for deep FL imaging. From the commercially-available VSD compound (IR780 perchlorate), 8.30%, 49.41%, 69.69%, and 80.95% of fractional contrasts in FL emission would be achievable with 1, 3, 6, 9-μM VSD concentrations (Figure S1).^18,19^ Our team also presented the feasibility of transcranial FL VSD neuroimaging using this VSD compound.^21^

In this paper, we present an *in vivo* proof-of-concept of functional intra-operative FL localization of the near-infrared cyanine VSD responses evoked by penile erection stimulation using a rodent animal model. Our major hypothesis is that the FL quenching-based VSD mechanism presents the functional contrast of the periprostatic erectogenic nerve network that reacts to an electrical stimulus in real-time. The temporal response time for VSD redistribution occurs at a sub-second scale^18^, suggesting that the real-time calibration for a surgeon’s performance will be enabled during RP. Also, we applied direct VSD administration to the rat periprostatic nerves situated within intact levator and periprostatic fascia layers. The staining time is designed to be within 10 minutes so that it would not delay or hamper the current standard RP protocol.

## Results

### Functional FL imaging performance characterization

Figure 1 presents our intra-operative FL imaging system for erectogenic nerve network localization *in vivo*. The spatial resolution of the imaging fiberscope was evaluated using a USAF 1951 Resolution Target (R3L3S1P, Thorlabs, New Jersey, USA) with line pair designed in multiple contrast Group and Element (Figure 2a). The best resolution of the system along a horizontal line (Y ROI) and vertical line (X ROI) is within Group 2, Element 1 and 2, indicating the system spatial resolution is within 111.36 μm to 125 μm. The minimal sensitivity threshold in VSD concentration was also estimated. The VSD droplets at 0.1, 0.25, 0.5, 1, and 4mM concentrations were imaged by the same imaging system, and the measured FL intensities were fitted into an exponential curve using MATLAB software (R^2^: 1; Root mean square error: 16.14) (Figure 2b). As presented in the magnified plot, the minimal sensitivity threshold was estimated at 6.77μM in VSD concentration, by having a cross-point between fitted curve and a noise level measured without laser excitation (i.e., 361 in arbitrary FL intensity).

**Figure 1.**
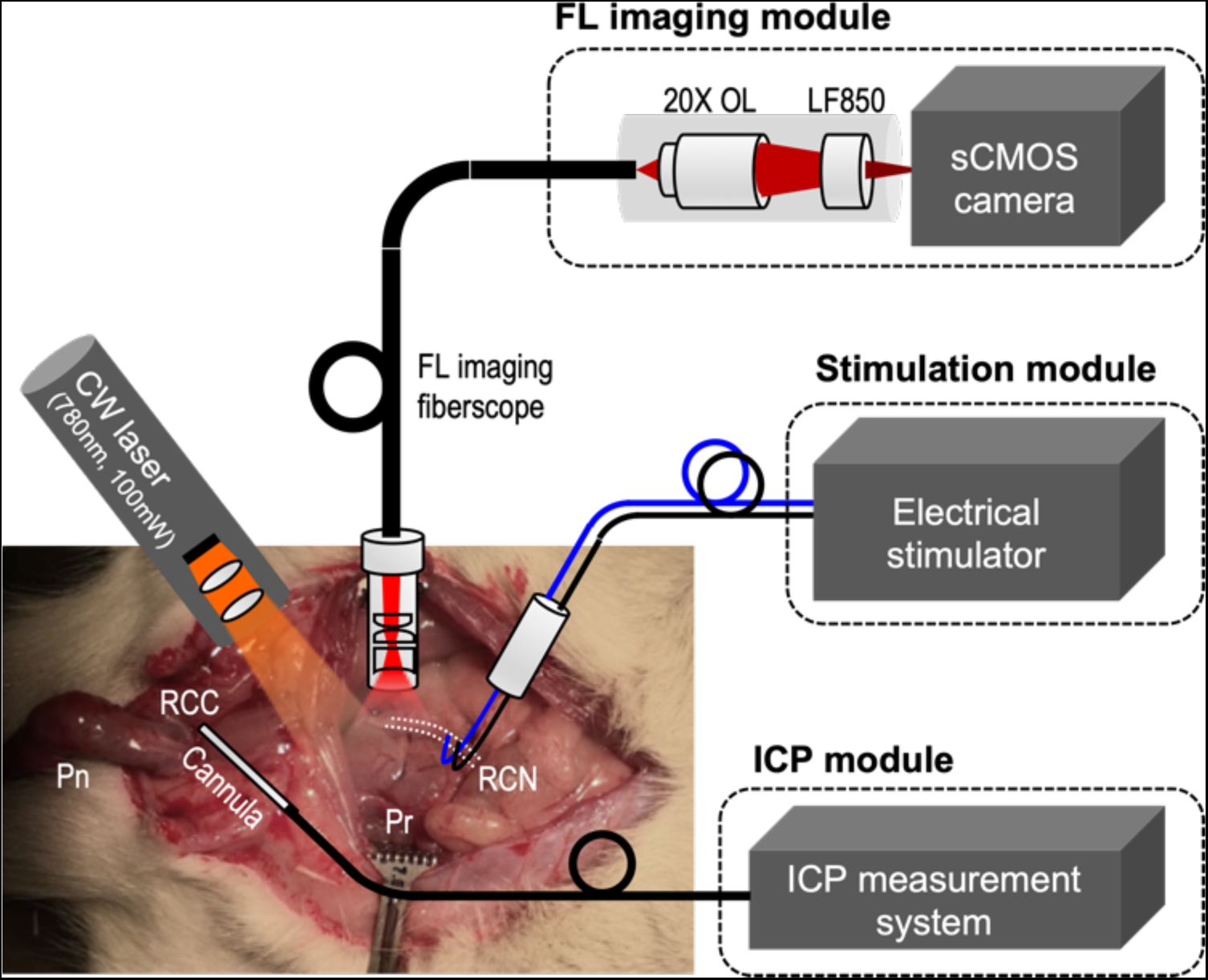
Intra-operative fluorescence (FL) imaging of rat prostatic nerve network *in vivo*. OL, objective lens; LF, emission longpass filter. Pn, penis; RCC, right corpus caverosum; Pr, prostate; RCN, right cavernous nerve; ICP, intracavernosal pressure.

**Figure 2.**
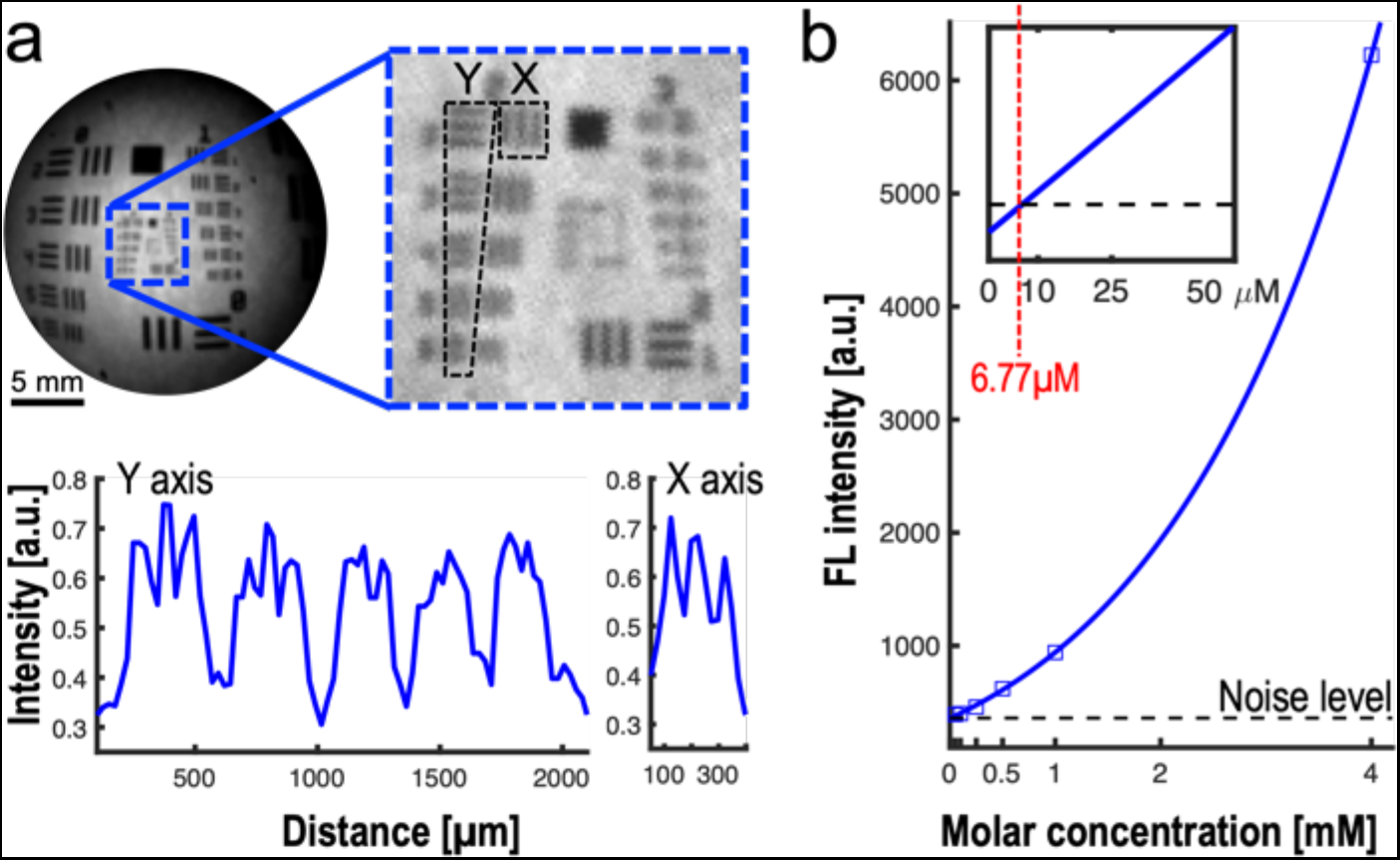
Functional intra-operative FL imaging system characterization. (a) Spatial resolution assessment of the fiberscope using a USAF 1951 target. (b) System sensitivity curve estimated from FL imaging of different VSD solution at 0.1, 0.25, 0.5, 1, and 4mM concentrations. The magnified plot in the upper-left side indicates the cross-point at 6.77μM between fitted sensitivity curve and noise level without laser excitation (dotted black line).

### Characterization of temporal VSD response

Figure 3 presents temporal characterization of the VSD responses using lipid vesicle model, mimicking neuronal membrane depolarization from polarization at around −120 mV. We used a framework already established in our previous studies.^18,19^ The first phase change induces the membrane transition from neutral state to polarized state, and triggers transmembrane VSD influx into the lipid vesicle. This VSD redistribution resulted in FL quenching at −7.51±2.22 %. The transition time of the VSD responses slowly took effect: 1.80 seconds and 4.85 seconds for 50 % and 90 % changes, respectively. On the other hand, the second phase change with gramicidin triggers membrane depolarization and corresponding VSD dispersion to both internal and external spaces of the lipid vesicle. This process presented up to 16.21 % increase in FL intensity within sub-second transition time: 128.53 ms and 192.79 ms for 50 % and 90 % changes, respectively, suggesting FL dequenching effect with near-infrared VSD compounds.

**Figure 3.**
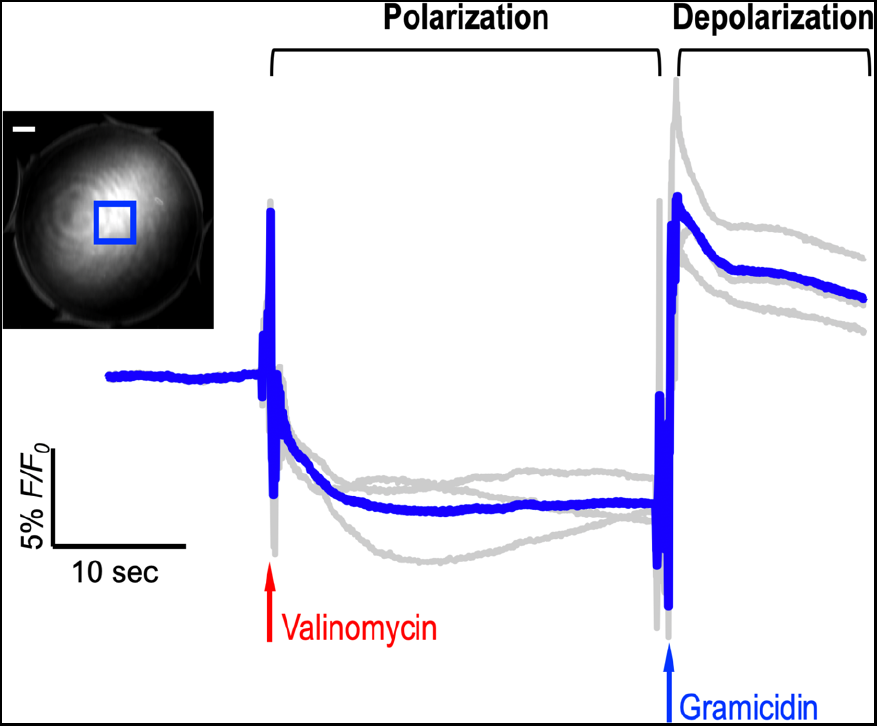
Temporal characterization of VSD response using lipid vesicle membrane model. Gray data indicate individual FL trace measurements. Each trace was normalized with the FL intensity before the administration of valinomycin. Initial signal fluctuations in polarization and depolarization phases are due to the administration of valinomycin and gramicidin. Blue rectangular in the exemplary FL image indicates region-of-interest, and white bar shows 1 mm. Gray lines were from three trials. Note that the FL contrast change obtained from the setup would be different from our previous study with a calibrated spectrofluorometry^18,19^, as only partial depth in a droplet would react to the ionophores manually administrated with pipette.

### Intracavernosal pressure monitoring

Intracavernosal pressure (ICP) measurement was performed for two reasons; 1- To validate that the nerve localized with the VSD is the CN, and 2- To determine whether the VSD has damaging effect on the cavernous nerve. Figure 4a shows the experimental design for ICP measurement induced by CN electrical stimulation in vivo. The significant ICP increase induced by electrical stimulation of the CN was measured following VSD staining: basal and maximal ICP values in VSD group (n=3) were 15.14±2.93 mmHg and 70.62±15.18 mmHg, respectively. Similarly, as expected, the control group without VSD staining (n=3) presented comparable level of ICP tracing with basal and maximal ICP values of 9.22±3.10 mmHg and 69.89±1.92 mmHg, respectively. There was no statistical significance between VSD and control groups (p = 0.07 and 0.94 for basal and maximal ICP values, respectively) (Figure 4b). The stimulated erectile response in the VSD group was also validated with the observations of tumescence and detumescence, as shown in Figure 4c and Movie 1. The results indicate that, within our proposed experimental paradigm, erectile function was preserved without damaging effect on the CN following VSD administration at given concentrations, staining durations, and flushing procedure.

**Figure 4.**
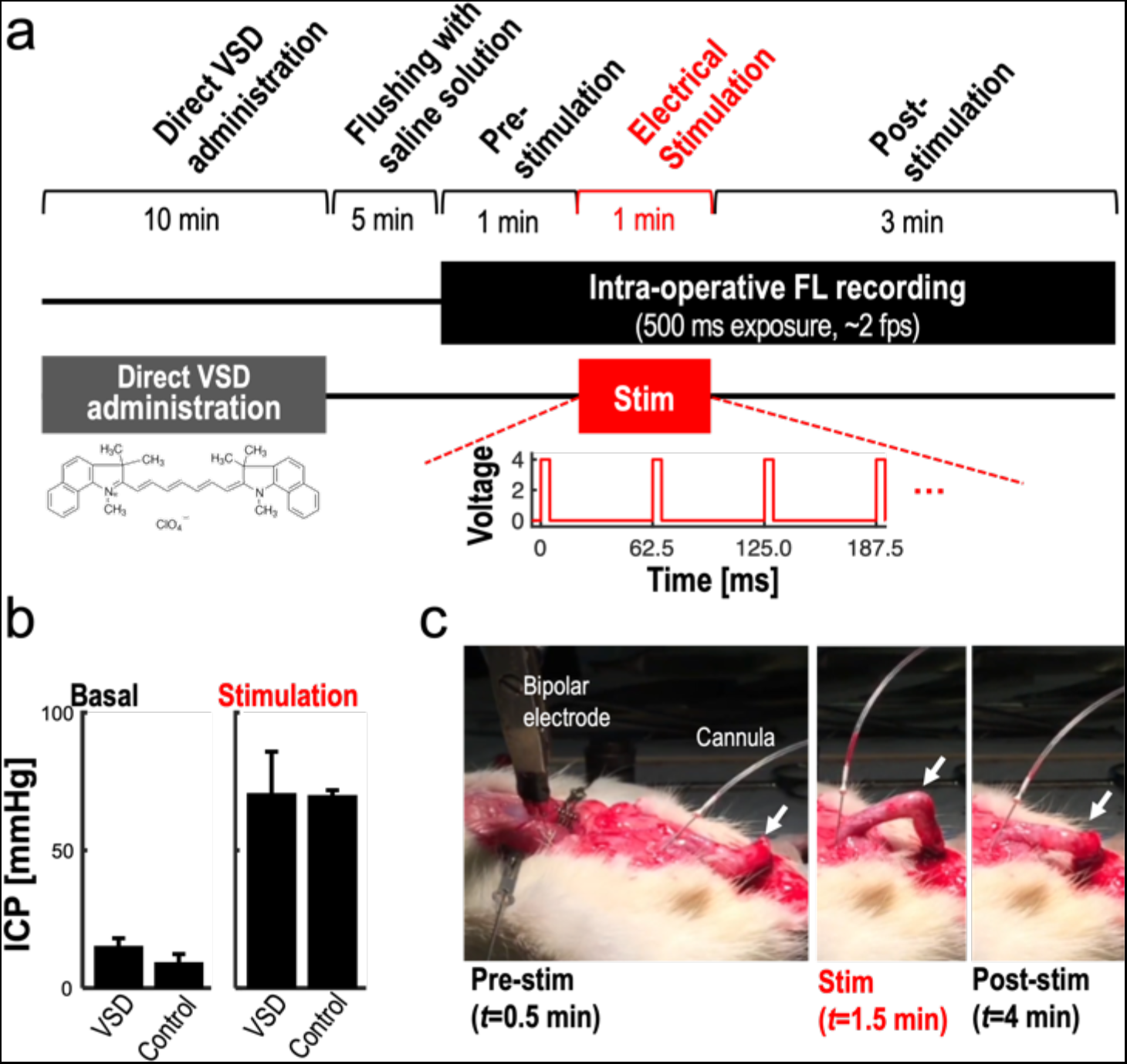
*In vivo* experimental setup for a proof-of-concept of prostatic nerve imaging using near-infrared cyanine VSD: (a) *in vivo* experimental protocol. (b) The validation of no adverse effect of the direct VSD administration using intracavernosal pressure (ICP) measurements before and after the electrical stimulation in control and VSD groups. (c) Corresponding observation of erectile function at each experimental phase (Movie 1).

### Intra-operative nerve localization

Figure 5a presents the white-light and FL images obtained from the given FOV on the rat prostate. The right CN (RCN) branching from the major pelvic ganglion (MPG) was differentiable by the naked eye during the surgery in by the white-light imaging, and apparent FL emission was detected from the entire prostate surface after performing direct VSD administration and flushing out procedures. This result confirmed the successful trans-fascial penetration of VSD. Figure 5b shows the time-averaged evolutions of fractional FL intensity change (*F*/*F*_0_) at each image pixel in pre-stimulation, stimulation, and post-stimulation phases from the reference frame averaged for 0 – 0.5-minute duration in pre-stimulation phase (across 60 frames). The stimulation phase revealed respectively up to 10.56±4.14 % and 7.04±4.77 % of *F*/*F*_0_ at the CN and CNB structures with photo-bleaching correction (Figure S3). These results may represent the chain reaction of the nerve network from the CN to CNB with different nerve areas under stimulation. Note that the bright spot in the pre-stimulation phase is due to the reflection of the time-variant fluctuations of the excitation laser. Our emission filter can reject most of the reflection, but there could be such a residual. However, there was no reflection in the prostate region-of-interest by having an angle between excitation and FL recording (Figure 1). The real-time video is shown in Movie 2.

**Figure 5.**
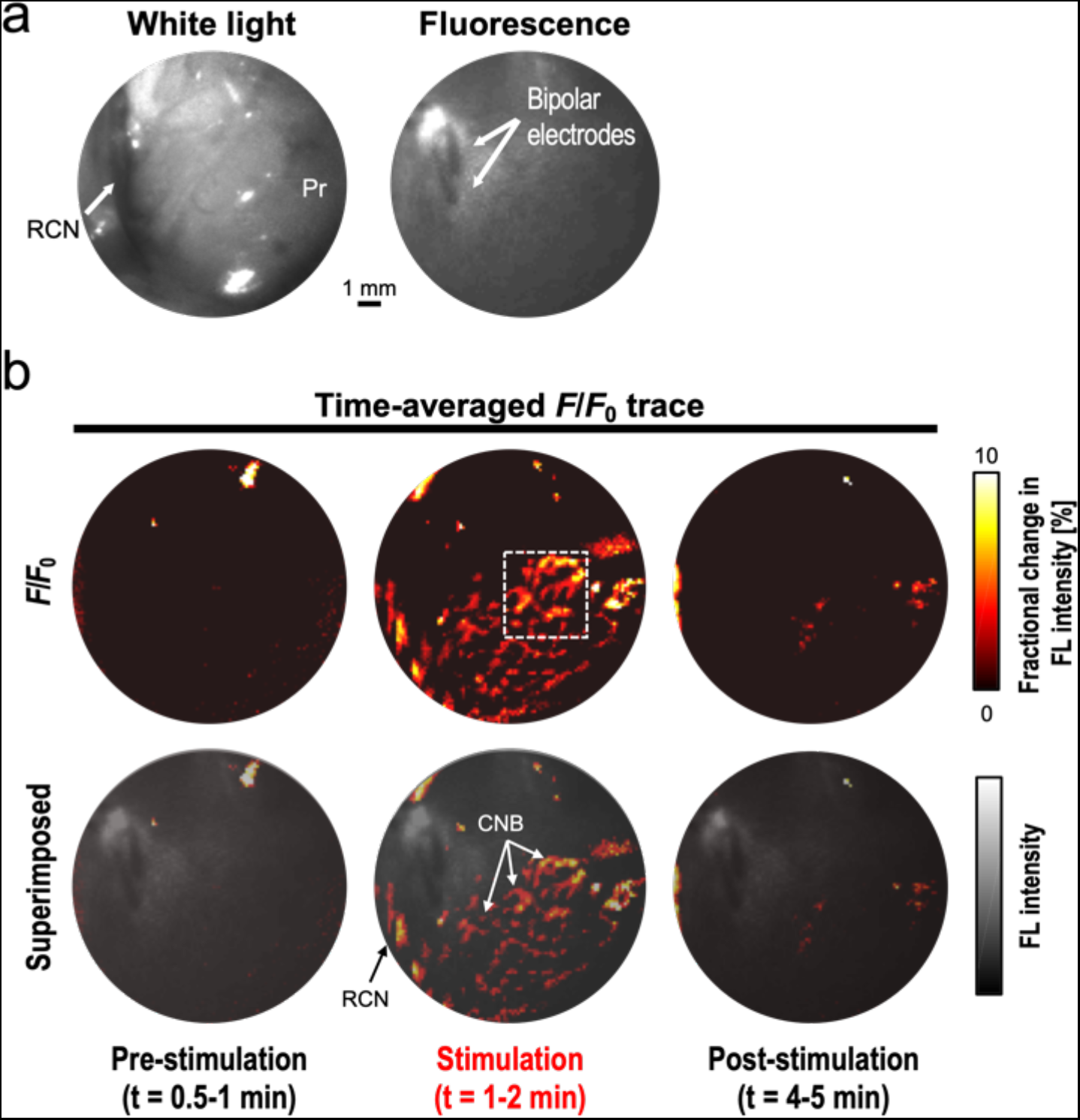
*In vivo* intra-operative nerve localization: (a) white-light and fluorescence (FL) imaging, (b) time-averaged *F*/*F*_0_ images during stimulation and pre-/post-stimulation phases. RCN, right cavernous nerve; Pr, prostate. The real-time movie of *F*/*F*_0_ image sequence is available in Movie 2.

### Evaluation of motion artifacts during electrical stimulation

The slight motion in the region-of-interest (ROI) was generated conceivably by the instantaneous blood volume change as a product of electrical stimulation, but it was not significant in our validation study. The normalized cross-correlation coefficient was calculated from the 4×4-mm^2^ region-of-interest (ROI) indicated by the dotted square in Figure 5b. Note that this ROI was selected as it yielded the highest *F*/*F*_0_ from the prostate surface. The normalized cross-correlation has calculated from the reference FL image averaged during the first half minute in the pre-stimulation phase (0 – 0.5 minute, 60 frames). From this, the worst correlation coefficient was at 0.979 during the stimulation and post-stimulation phase (Figure 6a). This confirms the fractional FL contrast in the ROI was not generated by any structural change over time. Also, we made a counter-hypothesis that the mean FL intensity in the ROI should be constant regardless of stimulation if the contrast was adversely caused by motion artifacts. For this, the *F*/*F*_0_ change over the ROI was averaged at each *F*/*F*_0_ image. However, the global FL intensity in the ROI was increased in the stimulation phase and restored to the basal level in the post-stimulation phase (Figure 6b). This manner also confirmed that the fractional contrast was contributed by the VSD redistribution mechanism.

**Figure 6.**
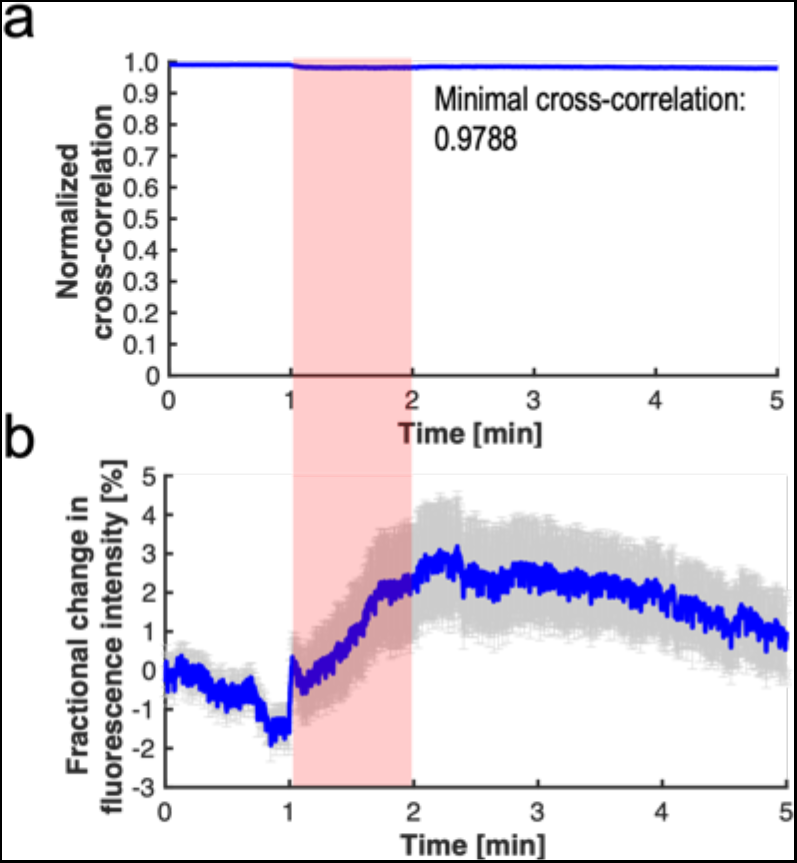
Evaluation of motion artifacts during *in vivo* experiment. (a) Normalized cross-correlation coefficient. (b) Fractional change in FL intensity over time at the entire region-of-interest.

### Histopathological validation of direct VSD administration

The direct VSD staining procedures were validated to localize rat nerves layered within the periprostatic and levator fascias. Frozen-sectioning and histopathological analysis were performed to demonstrate the penetration of VSD into the CN network in line with the direct administration protocol used. Figure 7 presents the white-light and FL microscopic images obtained from prostate slices, showing sufficient VSD staining depth into nerves interposed between the prostate gland and levator fascia whose thickness is a few hundred μm. The clear round cross-sections of nerve branches were successfully differentiated.

**Figure 7.**
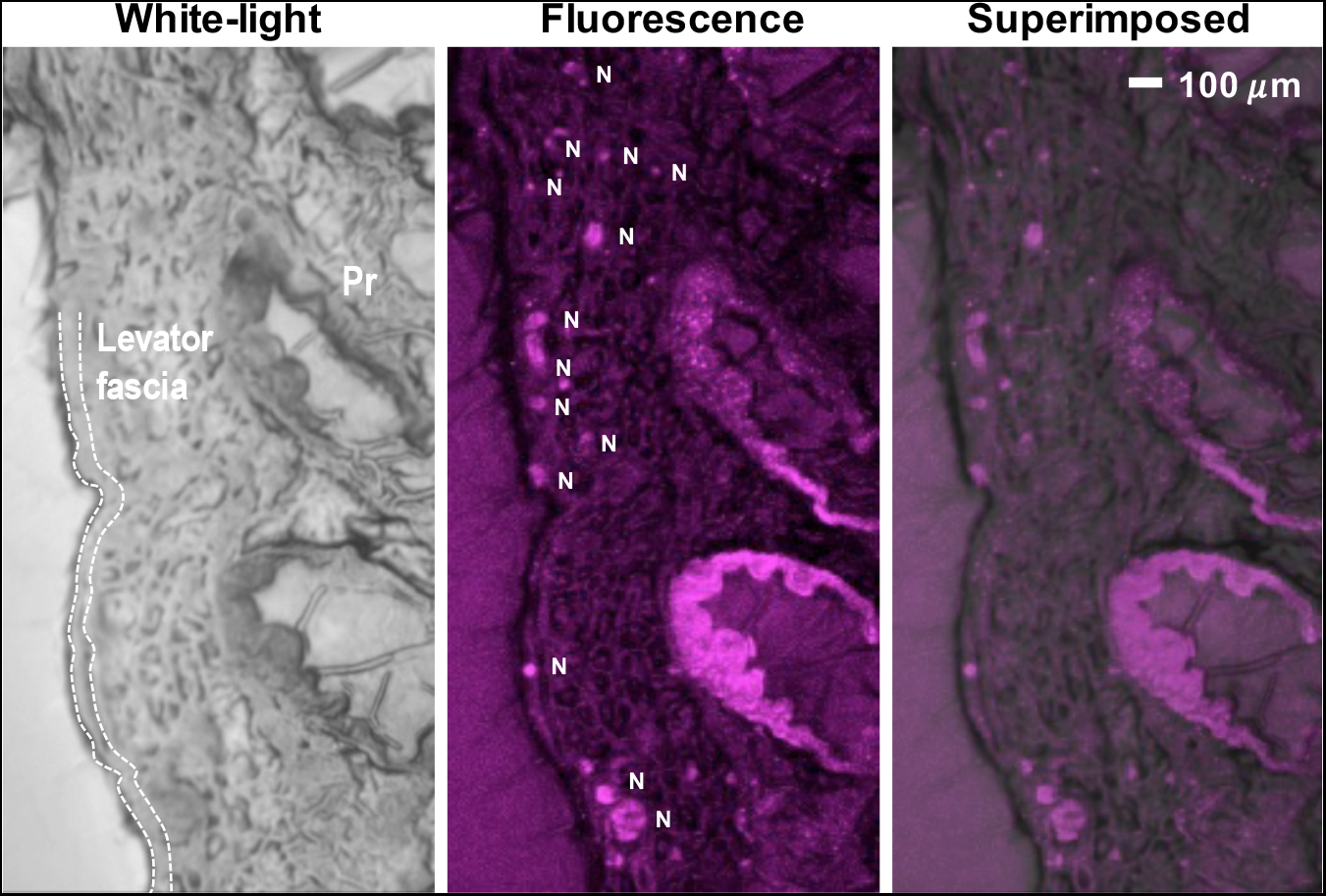
frozen-sectioning histopathological analysis on *ex vivo* rat prostate sample. Dotted line presents the levator fascia covering the prostatic fascia with CNB. White bar indicates 100 μm. Pr, prostate; N, nerve.

## Discussion

The prostate is understood to be the leading organ for new cancer cases and resulting deaths for males in the United States, (180,890, 21% in total cancer diagnosis), according to a 2016 estimate by Siegel, *et al.*^23^ Although survival rates have been improved by RP for patients with early-detected prostate cancer, a challenging assignment remains to prevent post-operative erectile dysfunction and/or urinary incontinence, which significantly impact quality of life. In this paper, we present an *in vivo* proof-of-concept of a novel functional nerve localization method that can be easily be amenable to the current standard RP protocol. The FL imaging system was first implemented, providing 6.77 μM of the minimal sensitivity threshold in VSD concentration, with 111.36 μm and 125 μm of spatial resolutions in horizontal and vertical directions, respectively (Figure 2). From the lipid vesicle membrane model, mimicking neuronal polarization and depolarization, the 50-% transition time of VSD response were measured at 128.53 ms and 1.80 seconds for membrane depolarization and polarization, respectively (Figure 3). Note that the VSD redistribution mechanism is usually slower than other voltage sensing mechanisms, but it can yield higher FL contrast.^24^ The functional FL imaging of the VSD-stained rat prostate presented the complex nerve network (CN + CNB) distinct from the surrounding tissues only when CN electrical stimulation was induced (Figure 5). This result was enabled by the trans-membrane VSD redistribution mechanism, which reflects depolarization events happening in the relevant nerve network in response to stimulation applied to the CN. Successful direct VSD administration to nerves was confirmed by histopathological analysis (Figure 7). Moreover, no adverse effects from direct VSD administration was found in ICP measurements (Figure 4). The 10-min duration for VSD staining will be applied between RP surgical steps and will not interrupt the procedure – Once surgeons access the prostate surface after cleaning the peri-prostatic tissue and endopelvic fascia incision, they will first apply the VSD and then perform the dorsal vein ligation during the VSD staining period which takes approximately 10 minutes. While we present these results in the open retropubic radical prostatectomy format on rodent animal, the proposed real-time, functional localization of cavernous nerve network should also be effective in laparoscopic configurations. Applying a digital FL imaging component for surgical guidance is becoming more common, and the direct VSD administration can be readily conducted through abdominal ports. In all, we presented FL VSD imaging can yield functional specificity representing the responsivity of CN network intra-operatively, which cannot be given in existing affinity-based dyes. For example, during RP, surgeon may rapidly evaluate the wide surgical field when planning nerve-sparing incisions, and can be assisted to design the optimal surgical procedures for better clinical outcome. Also, on-site surgical evaluation could be performed right after all the nerve-sparing procedure. In this case, any delay or decrease in VSD signal transition may indicate the damage on CN network. This information would be very important to plan following-up treatment for erectile dysfunction and urinary incontinence.

We plan to produce further advances in VSD technology. The chemical characteristics of the VSD could be optimized to produce a higher fractional contrast change in FL emission between polarized and depolarized states while providing a higher absorption coefficient at the near-infrared range for better imaging sensitivity. Additionally, exploiting the positively-charged electrochemical property of the VSD may enhance staining efficiency based on increased responsiveness to the membrane potential variation in the CN nerve network. Furthermore, the temporality of the VSD redistribution may be an important parameter to more rigorously characterize. Theoretically, the repetitive trans-membrane VSD redistribution mechanism should be achieved in a sequence of depolarization and repolarization events. However, the results shown in Figure 3 and Figure S3 suggest the different temporal features of VSD redistribution in membrane depolarization and repolarization events during transitions between stimulation and post-stimulation phases, respectively. Therefore, further analysis of the VSD characteristics will enhance perspectives to optimize the real-time quantification of nerve functionality in real time. A correlation of functional FL imaging to a multi-electrode recording of erectile electrophysiology would yield convincing validation regarding how much membrane potential variation can be detected by the proposed FL VSD imaging. On the other hand, a direct comparison between affinity-based dyes and our VSD may give informative insights on the practical pros and cons of each method. We will also address toxicity concerns of the proposed cyanine VSD. Our team is optimistic of its safety because as it is composed of a chromophore identical to that present in FDA-approved indocyanine green (ICG). In the proposed method, the direct tissue contact of the VSD solution will be limited to the prostatic region for 10 minutes at most, and the amount perfused into a patient’s bloodstream will be negligible. For these reasons, the possibility of chemical toxicity effects for a patient’s body is likely low. We already presented that the direct VSD administration did not exert any adverse objective or subjective effects on erectile function (Figure 4b,c). We will validate this observation in future investigations.

The detailed methodology of the direct VSD staining protocol could be improved to represent a more quantitative measure of membrane potential variations. The local VSD concentration in nerve network will be dependent to the permeability of the fascia layers and the disposed VSD concentration on the surface when a sufficient VSD volume was applied. However, in the *in vivo* results presented in this paper, the majority of the VSD solution ran down from the prostate surface based on the angled position, and only the superior portion of the prostate accumulated the VSD solution. This condition possibly led to the results in Figure 5b producing the highest *F*/*F*_0_ at the center of the prostatic region. Therefore, optimizing the direct trans-fascia VSD delivery will be pursued in our future studies. A patch-based delivery method may well be investigated to achieve both fast, uniform VSD delivery, providing quantitative measures in membrane potential variation at the erectogenic nerve network.

In our future studies, a larger number of rodent animals will be included with the improved technologies and experimental setup, and the practical CNB contribution to erectile function will be studied in greater detail. The extent of the localized nerve damage induced to CNB with various distances from the primary CN will be assessed, and the area of the damage will be controlled based on various techniques.^25^ CN stimulation and ICP measurement protocols will also be applied for quantitative evaluation. This investigation will show the extent to which part of the CNB contribute to the erectile function in addition to the primary CN.

We are very eager to extend this concept to a multi-modal surgical guidance approach incorporating photoacoustic (PA) and FL imaging modalities. There have been several extensions into multi-modal imaging for intra-operative guidance: MRI + transrectal US^26^; PA microscopy (PAM) + OCT^27^; PET + FL^28^; US + PA tomography (PAT) + FL^29,30^. However, no such attempts can achieve enough synergy without a multi-modal VSD indicator for any proposed application. PA imaging is an emerging hybrid imaging modality providing both optical absorptive contrast and acoustic imaging depth, and it also supports versatile imaging scales by microscopic (5 μm of spatial resolution for up to 2 mm-depth) and tomographic imaging configurations (~800 μm of spatial resolution for up to 7-cm depth).^31–33^ Also, recently, several PA neuroimaging researchers have proposed to quantify membrane potential variations using a genetically-encoded calcium indicator or non-radiative voltage sensor.^34,35^ We have also shown the concept of novel functional PA imaging of membrane potential variation *in vivo* using the same cyanine VSD used in this study. The study was based on the complementary PA and FL contrast, according to the stated VSD redistribution mechanism. We have already validated the VSD mechanism via controlled lipid vesicle experiments with various membrane potential levels and further presented *in vivo* brain imaging using rat seizure and NMDA infusion models.^19,36^ Based on these achievements, the multi-modal approach can be applied to nerve-sparing prostatectomy, which may promise more comprehensive guidance extended to a broad set of surgical procedures. For example, a pre-operative PA VSD imaging may delineate nerve distributions before making incisions; intra-operative guidance using reciprocal contrast in PA and FL imaging may maximize sensitivity based on membrane potential; post-operative monitoring using non-invasive transrectal PA imaging may assess erectile function recovery. We also developed a dual-modal imaging system using a pulsed Nd:YAG optical parametric oscillator (OPO) laser, which produces FL and PA contrast from the same absorber target. Further investigation will lead to optimized engineering solutions to facilitate its translation into pre-clinical animal studies and, eventually, to human investigations. Some surgeons may prefer to keep the constant nerve distribution map during nerve-sparing procedures, along with the functional, multi-modal contrast. Simple image fusion of the snapshotted VSD contrast will be developed to provide the nerve distribution map during a surgery if needed.

A more clinically relevant model will be considered for our future validation studies. Even though the current experimental results are encouraging with the rodent animal model, the guidance specification may not be optimal for larger animal models or humans – In the rat prostate, the fascia layers surrounding the prostate are very thin at around 100 μm of thickness, whereas the human prostatic fascia is much thicker. In practice, variability in fascia thickness should be evaluated because this factor may substantially affect the trans-fascial FL imaging and effectiveness of the VSD staining procedure. For this investigation, an *ex vivo* human prostate sample with intact levator and periprostatic fascia layers will be used. VSD droplets with various concentrations will be administered on the sample for various durations, and depths of penetration will be recorded with measurements of local sample thickness. FL imaging would be performed to validate the imaging depth, and the VSD-stained depth will be quantified by subsequent histopathological analysis. This study design would serve to optimize the VSD administration scheme as a function of fascia thickness for further translational *in vivo* evaluation using large animal models (e.g., canine, porcine, etc.) and/or a human trial. In the *in vivo* experiments, the instant calibration of the direct VSD administration setup will be allowed by a local fascia thickness given by pre-operative tomographic imaging with narrow axial resolution (e.g., US, OCT) or by pre-operative imaging (e.g., CT, MRI).

## Methods

### Near-infrared FL imaging of electrically-stimulated erectile function

The FL imaging fiberscope was designed to provide a similar endoscopic view as in a flexible clinical fiberscope. At the distal end of the fiberscope is a customized optical assembly to focus high angle beams (70-degree field of view) from the object into the fiber core body. The focused image at the distal fiber end is then relayed multiple times to the proximal fiber end through total internal reflectance. The customized optical assembly with 1.5-mm lens diameter was optimized using Optics Studio 15 SP1 (Zemax, Kirkland, Washington, USA), fabricated, and housed in front of the fiber relay body. The simulation layout, spot diagram, and modulated transfer function (MTF) of the optical assembly is shown in Figure S2. The resultant numerical aperture (NA) of the fiberscope was approximately 0.398. The fiber relay body (not shown in Figure S2) is a fiber bundle with 50,000 cores in 1,100-μm diameter at 4.5-μm pixel center-to-center spacing (FIGH-50-1100N, Fujikura Image Fiber Bundles, Myriad Fiber Imaging, MA). The relayed image near the proximal fiber end is focused on a scientific CMOS sensor (Hamamatsu ORCA-Flash4.0) using a 20X microscope objective lens (Bausch & Lomb Objective, Microscope Central, PA). The FL signal from the collected relayed image is filtered to the sensor with a long pass hard-coated dielectric coating filter at the cut-on wavelength at 850 nm (FELH0850, Thorlabs, New Jersey, USA). On the other hand, for the laser illumination, a 100-mW laser diode with central wavelength at 785 nm (FWHM 3 nm) equipped with a variable beam expansion lens is mounted on a separated arm to illuminate the sample. The position of laser illumination was aimed to cover a rat prostate region with illumination circle of 1 cm. The overall system configuration with the imaging fiber, the laser source, the nerve excitation, and ICP monitoring is shown in Figure 1.

### Voltage-sensitive dye preparation

The VSD (IR780 perchlorate, 576409, Sigma-Aldrich, Inc., United States) was vortexed in 5% DMSO, 5% Cremophor EL until dissolved, then dilute with 0.9% saline to be 1mM concentration for the proposed *in vivo* experiments. The peak absorbance and FL emission are at 790nm and 820nm, respectively.^18,19^

### Transient VSD response in lipid vesicle membrane model

The temporal VSD response was characterized using a lipid vesicle membrane model. Translucent lipid vesicle suspension was prepared as described in our previous literature.^18,19^ 25-mg soybean phosphatidyl- choline (type II) was first suspended in 1 mL of K^+^ buffer with 100 mM K_2_SO_4_ and 20 mM HEPES, and then vortexed for 10 min and sonicated for 60 min. VSD solution was diluted to 6 μM and added to the suspension. The temporal resolution of trans-membrane redistribution mechanism should be independent from the VSD concentration applied on lipid vesicle membrane. Adding 10 μL of the suspension to 1 mL of Na^+^ buffer (100 mM Na_2_SO_4_ and 20 mM HEPES) resulted in an approximately 100∶1 K^+^ gradient across membrane. The droplet of the lipid vesicle solution (100 μL) was place on a transparent 96-well plate cover for continuous FL imaging. From the setting, membrane polarization was induced by a K+ specific ionophore (2.5 μL of 10 μM valinomycin). On the other hand, a subsequent addition of nonspecific monovalent cation ionophore (2.5 μL of 1 mM gramicidin) induced membrane depolarization. During FL recording, valinomycin and gramicidin were injected with 30 sec latency. The photobleaching was corrected with 30-sec pre-injection phase duration. Note that the FL contrast change obtained from the setup would be different from our previous study with a calibrated spectrofluorometry ^18,19^, as only partial depth in a droplet would react to the ionophores manually administrated with pipette. The experiment was designed to address our primary interest, temporal characterization of VSD response.

### Animal preparation

Adult male Sprague-Dawley rats (325-350g; Charles River Breeding Laboratories, Wilmington, MA, USA) were used. All experiments were approved by the Johns Hopkins University School of Medicine Institutional Animal Care and Use Committee in accordance with the National Institutes of Health Guide for the Care and Use of Laboratory Animals. The rats were anesthetized with an intraperitoneal injection of a ketamine (100mg/kg) and xylazine (5mg/kg) mixture. The prostate was exposed via a midline abdominal incision, and CNs and MPGs were located bilaterally posterolateral to the prostate.

### *In vivo* experimental protocol

Figure 4a presents the *in vivo* experimental protocol employed in this study. The 200μl of 1mM VSD in DSMO + Cremophore EL solvent was directly administrated to the rat prostate surface for 10 minutes before starting FL recording. The VSD not bound to the prostate nerve membrane was flushed out for 5 minutes with 2-ml phosphate buffered saline (PBS) solution. For the prostate with VSD administration, FL images were recorded for 5 minutes, and 1 minute of stimulation was induced from 1 minute to 3 minutes. There were 1 minute of pre-stimulation and 3 minutes of post-stimulation phases to track the change of VSD response.

### Supramaximal electrical CN stimulation and intracavernosal pressure monitoring

For the continuous ICP monitoring, the shaft of the penis was denuded of skin and fascia, and the right crus was punctured with a 23-gauge needle connected via polyethylene-50 tubing to a pressure transducer. For electrically stimulated penile erections, a bipolar electrode attached to a Grass Instruments S48 stimulator (Quincy, MA, USA) was placed on CNs. Supramaximal stimulation was induced for 1 minute with 4 volts, 16Hz, and 5-msec square-wave duration.^22^ ICP was recorded (DI-190, Dataq Instruments, Akron, OH, USA) for 5 minutes; pre-stimulation (1 minute), stimulation (1 minute), and post-stimulation (3 minutes). Results were analyzed using the MATLAB program (Mathworks, Natick, MA, USA). At the conclusion of experiments, animals were sacrificed by a lethal intracardiac injection of saturated potassium chloride and prostate, both MPGs, and CNs were collected.

### Image acquisition and reconstruction

The frame rate of the CMOS camera was approximately 2fps, with 500 msec of exposure time. The FL response recorded from the rat prostate was processed in an offline workstation. The FL frames recorded during the first half minute (0 minute – 0.5 minute) without any electrical stimulation were averaged to form a reference image. Afterward, the fractional change of FL emission (*F/F*_0_) was calculated at each pixel and at each time point to derive the VSD response evoked by electrical stimulation.

### Frozen-section histopathological analysis of *ex vivo* rat prostate samples

After getting the FL recording, a whole rat prostate was immediately harvested. It was placed in fresh 10% formalin for more than 48 hours with gentle agitation using a conical rotator. Cryoprotection processing was done via a series of sucrose gradients (15%, 20%, 30% for 12-24 hours each). Prostates were frozen-sectioned at 300μm thickness. Slides with tissue sections in ProLong Diamond Anti-face mountant were imaged using the LI-COR Odyssey for FL visualization of VSD perfusion.

## Acknowledgments

Financial support was provided by the NIH Brain Initiative under Grant No. R24MH106083-03, the NIH National Institute of Biomedical Imaging and Bioengineering under Grant No. R01EB01963, the NIH National Institute of Diabetes and Digestive and Kidney Diseases under Grant No. R56DK114095, and NIH National Institute of Heart, Lung and Blood under grant number R01HL139543. Jeeun Kang, Ph.D. is supported by Basic Science Research Program through the National Research Foundation of Korea (NRF) funded by the Ministry of Education (2018R1A6A3A03011551) and Early Investigator Research Award, Congressionally Directed Medical Research Programs, U.S. Department of Defense (DOD), United States (W81XWH-18-1-0188). Serkan Karakus, M.D., is supported by grant from the National Institutes of Health, USA (NIH/NIDDK grant R56DK114095 to A.L.B.).

## Author contributions statement

J.K. designed, delineated, and performed *in vivo* experiments, data processing and analysis, and wrote the first manuscript. H.N.D.L. customized the FL imaging module for intra-operative applications and applied to *in vivo* experiments. S.K. prepared the animal model and conducted *in vivo* experiments. A.M. conducted a histopathological validation of the direct VSD administration under supervision by M.M.H. E.M.B. conceived the research, and J.U.K., A.L.B., and E.M.B. co-directed the project. All authors edited the manuscript.

