## Supplementary Information for "Real-time, functional intra-operative localization of rat cavernous nerve network using near-infrared cyanine voltage-sensitive dye imaging"

**Spectrophotometric/fluorometric characteristics of FL quenching-based cyanine VSD.** In the spectrophotometric-/fluorometric measurements (Figure S1a), spectral peaks were at 790nm and 820nm for absorbance and FL emission, respectively. We already tested a VSD with the comparable chemical structure (PAVSD800-2).^18^ The testing was conducted using a lipid-vesicle membrane model with different VSD concentrations: 1, 3, 6, 9μM. When adjusting the FL contrast changes, 8.30%, 49.41%, 69.69%, and 80.95% of fractional contrasts in FL emission would be achievable at 1, 3, 6, 9-µM VSD concentrations, respectively.

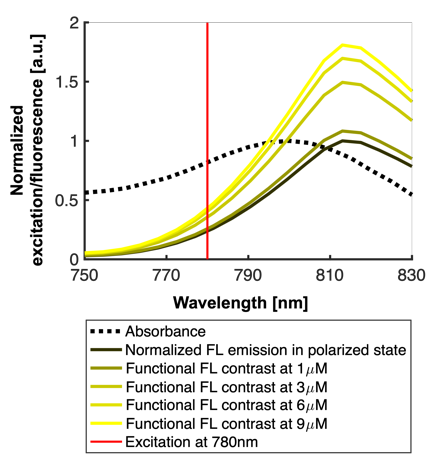

**Figure S1.** Spectrophotometric/fluorometric measurement for VSD solution. Red line indicates the excitation wavelength used in our *in vivo* experiments, i.e., 780nm.

**Customization of optical assembly.** The customized optical assembly is a collection of lenses to collect high angle beam and form an image at the fiber bundle surface for further image relay (Figure S2). The optical assembly is designed and optimized using Optics Studio 15 SP1 (Zemax, Kirkland, Washington, USA) with the known parameters of viewing angle within 60-70 degree and the lens diameter is within 1.5 mm. The fiber NA is approximately 0.398. Figure S2 shows the simulation layout, spot diagram, and modulated transfer function (MTF) of the customized optical assembly.

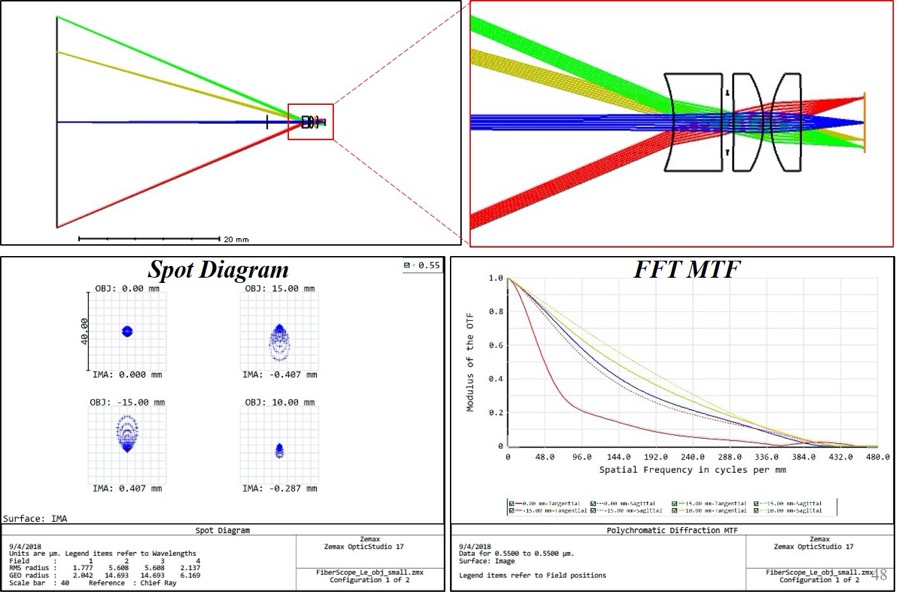

**Figure S2.** Optical design of the focusing assembly for the fiberscope.

**Localized analysis of fractional FL intensity changes in nerve regions.** The localized analysis in nerve branches in FL image was performed. In the FL image sequence, ROIs presenting CN, and CNB were selected as shown in Figure S4 with dotted rectangular enclosures. On the other hand, the background (BG) region was also selected to correct the photo-bleaching presented over time (rectangular enclosures with solid line). The appropriate selection of BG region was confirmed in the pre-stimulation phase with nulled fractional change in FL intensity. Note that the correction could not be applied in entire FOV as the amount of photo-bleaching is varied in spatial domain.

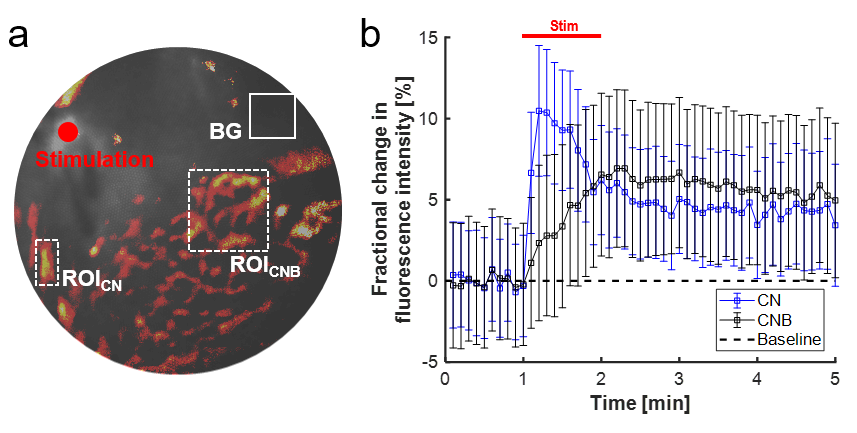

**Figure S3.** Fractional FL intensity change on localized nerve regions: (a) locations of regions-of-interest (ROIs). BG, background; ROI_CN_, CN region; ROI_CNB_, CNB region; (b) the plot of fractional change in FL intensity at each ROI. Note that the plots were corrected by the FL intensity trace quantified at the BG region.
